# Biodiversity shows unique responses to land-use change across regional biomes

**DOI:** 10.1101/2023.03.08.531730

**Authors:** Peggy Bevan, Guilherme Braga Ferreira, Daniel J Ingram, Marcus Rowcliffe, Lucy Young, Robin Freeman, Kate E. Jones

**Affiliations:** Centre for Biodiversity and Environment Research, University College London, Gower Street, WC1E 6BT; Institute of Zoology, Zoological Society of London, Wellcome Building, Outer Circle, London NW8 7LS; Durrell Institute of Conservation and Ecology, School of Anthropology and Conservation, University of Kent, Canterbury, CT2 7NR, UK; WWF UK, Living Planet Centre, Brewery Road, Woking, GU21 4LL

## Abstract

Biogeography has a critical influence on how ecological communities respond to threats and how effective conservation interventions are designed. For example, the resilience of ecological communities is linked to environmental and climatic features, and the nature of threats impacting ecosystems also varies geographically. Understanding community-level threat responses may be most accurate at fine spatial scales, however collecting detailed ecological data at such a high resolution would be prohibitively resource intensive. In this study, we aim to find the spatial scale that could best capture variation in community-level threat responses whilst keeping data collection requirements feasible. Using a database of biodiversity records with extensive global coverage, we modelled species richness and total abundance (the responses) across land-use types (reflecting threats), considering three different spatial scales: biomes, biogeographical realms, and *regional biomes* (the interaction between realm and biome). We then modelled data from three highly sampled biomes separately to ask how responses to threat differ between regional biomes and taxonomic group. We found strong support for regional biomes in explaining variation in species richness and total abundance compared to biomes or realms alone. Our biome case studies demonstrate that there is a high variation in magnitude and direction of threat responses across both regional biomes and taxonomic group, but all groups in tropical forest showed a consistently negative response, whilst many taxon-regional biome groups showed no clear response to threat in temperate forest and tropical grassland. Our results suggest that the taxon-regional biome unit has potential as a reasonable spatial and ecological scale for understanding how ecological communities respond to threats and designing effective conservation interventions to bend the curve on biodiversity loss.

## Introduction

Despite multiple internationally agreed science-based conservation targets, biodiversity continues to decline across the globe, with the consequent loss of essential ecosystem services (Tittensor *et al*., 2014; Brooks *et al*., 2015; Ceballos, Ehrlich and Dirzo, 2017; Isbell *et al*., 2017; Mace *et al*., 2018; IPBES, 2019; Ingram *et al*., 2021). Habitat disturbance caused by land-use change is a major factor contributing to biodiversity loss (Newbold *et al*., 2015). However, the impact of disturbance on biodiversity (its response) varies spatially and is contingent on factors such as the intensity of environmental change (Felipe-Lucia *et al*., 2020), the original habitat (Monsarrat, Jarvie and Svenning, 2019) and taxon studied. Such threat-response relationships will further be influenced by specific life-history traits, biogeographic context, and the ecological scale being measured (*e*.*g*., a community, population or individual) (Isaac and Cowlishaw, 2004; Murphy, 2021; Suraci *et al*., 2021). Given the recognised variability of biodiversity’s responses to threats such as habitat disturbance, this knowledge can help improve conservation decisions. With the adoption of new targets such as the Kunming-Montreal Global Biodiversity Framework (CBD, 2022) and the Global Deal for Nature (Dinerstein *et al*., 2019), it is vital to develop biodiversity monitoring systems that can accurately measure indicators at an appropriate spatial and ecological scale that optimises available resources with the resolution necessary to inform successful conservation actions.

The specific location of a species or ecological community on the planet, known as its biogeography, plays a crucial role in determining its sensitivity to an anthropogenic threat. This could be down to the current climatic circumstances of the location, historical natural disturbances, or the history of human activity in the area. For example, studies on biodiversity in tropical areas have been shown to have higher levels of fragmentation-sensitive species, stronger negative responses to land-use change and faster declining abundance than comparable temperate areas (Betts *et al*., 2019; Newbold *et al*., 2020; WWF, 2022). This could be explained by the extinction filter hypothesis, which suggests that ecosystems with a history of disturbance will be more resilient in the present day due to the previous filtering of sensitive species (Balmford, 1996). Tropical biomes are considered to have a history of low natural disturbance and a stable climate, whereas temperate biomes have been subject to elevated levels of glaciation, widespread fires, and human-caused forest loss in the last 10,000 years (Betts *et al*., 2019). In addition, traits typically associated with tropical species (Stevens, 1989; Gaston, 2000; Brown, 2014), including habitat specialisation and small range size, have also been associated with stronger negative responses to land-use change (Newbold *et al*., 2013). The history of land use and environmental disturbance is not uniform across the globe and will further contribute to modern day responses to anthropogenic threats (Goldewijk *et al*., 2011; Yang *et al*., 2021). Incorporating an understanding of biogeography and historical knowledge of an environment could play a key role in global biodiversity monitoring.

Considering the influence of biogeography on threat-response relationships, monitoring biodiversity at the smallest possible scale would be appropriate. However, there are currently more than 42,000 threatened species on the IUCN red list (IUCN, 2022), all of which cannot have unique conservation plans created for them. Therefore, indicators must be used at a higher ecological and spatial scale that strikes a balance between presenting global trends and understanding local scale responses to human pressure (Ingram *et al*., 2021). One way to account for biogeographic variation in threat responses is to use a monitoring framework based on spatial units, but choosing the correct unit is challenging. For example, creating national statistics for biodiversity is helpful for feeding into policy, but biodiversity is not nationally constrained (Murphy, 2021). A global framework has been suggested for monitoring biodiversity at an ecoregional level (Dinerstein *et al*., 2017, 2019; Smith *et al*., 2018). This framework is powerful when measuring variables that can be obtained using remote sensing or global species lists (*e*.*g*., Dinerstein *et al*., 2017; Smith *et al*., 2020), but many ecoregions are lacking the required field data or access to study population responses to human pressure at this resolution (Stephenson *et al*., 2015). It is essential to investigate whether broader spatial scales, already used in many global monitoring studies (Newbold *et al*., 2016; Blowes *et al*., 2019; WWF, 2022), can effectively encompass the diversity in threat-response relationships.

The broadest spatial unit of terrestrial habitats is the climatically defined biome, of which there are 14 (Olson *et al*., 2001). Responses of animal and plant populations to threats do differ between biomes (Greenville *et al*., 2018; Green *et al*., 2020) and species richness is particularly sensitive to land use change in tropical forest, tropical grassland and Mediterranean biomes compared to temperate and desert biomes (Newbold *et al*., 2020). In addition to biomes, there are 8 biogeographic realms, which broadly follow the continents (Olson *et al*., 2001; UNEP-WCMC 2004). Data from the Living Planet Database (Collen *et al*., 2009; WWF, 2022) has shown that decreases in vertebrate populations are more pronounced in southern hemisphere realms, particularly the Neotropics (Green *et al*., 2020; WWF, 2022). Although there is evidence for variation in threat-response relationships between biomes and between realms, the intersection between these two spatial units is rarely investigated, which may mean a large amount of variation is unaccounted for when monitoring biodiversity in biomes or realms alone.

A potential, intermediate spatial framework for monitoring biodiversity are regional biomes (the interaction of realms and biomes) (n = 64) (Ingram *et al*., 2021). Regional biomes each cover 11 ecoregions on average, ranging from 1-81 ecoregions (Olson and Dinerstein, 2002). Separating biomes by biogeographic realms can account for differences in evolutionary history, vegetative structure, threats, and socioeconomic status that occur between realms (Moncrieff, Bond and Higgins, 2016; Allan *et al*., 2019). Furthermore, threats are not spread evenly across the world and can be region-specific (Lewis, Edwards and Galbraith, 2015; Bowler *et al*., 2020); for example, tropical forest in southeast Asia is a threat ‘hotspot’, whereas large parts of the Amazon rainforest act as threat ‘refugia’ (Allan *et al*., 2019). Some areas of forest biomes, particularly in the northern hemisphere, are being afforested, whilst others experience mass deforestation (Song *et al*., 2018). In Asian tropical forests, species richness was found to be significantly more sensitive to disturbance compared to similar regions in South America and Africa (Gibson *et al*., 2011; Phillips, Newbold and Purvis, 2017). Conversely, vertebrate abundance is declining more in the Neotropics and Afrotropics than in the Indo-Pacific region (Green *et al*., 2020; WWF, 2022). These conflicting trends highlight the necessity of understanding and clearly reporting how biogeographical variation contributes to threat responses. Regional biomes may present a middle-ground between the finer scale monitoring of ecoregions and the coarser biome scale, facilitating robust reporting of biodiversity responses.

Here, we use the PREDICTS database to analyse data from over 400 field studies to explore the efficacy of regional biomes as a spatial monitoring unit for terrestrial species. Our expectation is that there will be negative responses to human-dominated land-use types compared to primary vegetation, but the magnitude of response will differ between regional biomes. We therefore hypothesise that regional biomes present a more parsimonious explanation of responses of biodiversity to land-use change than realms or biomes alone. Secondly, as case studies we model the three most data-rich biomes (Tropical Forest, Temperate Forest, Tropical Grasslands) in the PREDICTS database separately to explore how responses to land-use change and use-intensity vary between regional biomes. Here, we also consider the impact of taxonomic group (Vertebrate, Invertebrate, Plant) on the threat-response relationship within regional biomes. We expect the highest variation but also uncertainty in predicted responses between tropical regional biomes due to ecoregions being more distinct in these biomes (Smith *et al*., 2020), and smaller and less variable responses between temperate regional biomes due to the history of natural and anthropogenic disturbance leading to biotic homogenization (Newbold *et al*., 2018).

## Methods

### The PREDICTS database

We used species abundance and occupancy data from the PREDICTS database (Hudson et al 2017; 22678 sites, 480 sources, 666 studies, 47,000 species). We chose to use the PREDICTS database due to high global coverage compared to other databases (Hudson *et al*., 2017), making it suitable for a study comparing many regional biomes. The PREDICTS project (Predicting Responses of Ecological Diversity In Changing Terrestrial Systems; www.predicts.co.uk) is a database that collates biodiversity studies with comparable measures of terrestrial biodiversity from sites of different land uses and land-use intensity (Hudson *et al*., 2017). All sampled sites are classified by stage of disturbance or recovery which can be used as a substitute for actual temporal change at a single site (Walker *et al*., 2010; Srivathsa *et al*., 2018). This space-for-time substitution methods allows for specific measurements of biodiversity over distinct land-use types, acting as a proxy for anthropogenic pressure. PREDICTS does not include marine or freshwater data but does have extensive global coverage and has made attempts to reduce taxonomic and geographic bias (Hudson *et al*., 2017).

We summarised occurrence and abundance records by site and taxon using the predictsFunctions R package (Newbold, 2018). ‘Taxa’ were categorised as Vertebrate, Invertebrates, Plants or Fungi. The PREDICTS database covers 47 out of 64 regional biomes (70%), with some regional biomes more comprehensively sampled than others (Table S1). For analysis, we excluded data from three biomes with fewer than 50 sites in total from the dataset (Flooded Grasslands and Savannahs, Tundra, Deserts & Xeric Shrublands and Mangroves). Additionally, regional biomes were considered data deficient and excluded from analysis if they had no sites in primary vegetation land-use types (our reference level), or no sites in any kind of managed land use (cropland, pasture, and plantation forest). This ensured that regional biomes included in analysis had data points (sites) across a gradient of increasingly anthropogenic land-use types. After removing data deficient regional biomes, a total of 30 regional biomes were available for analysis (Fig. 1; 20011 sites, 545 studies).

**Figure 1.**
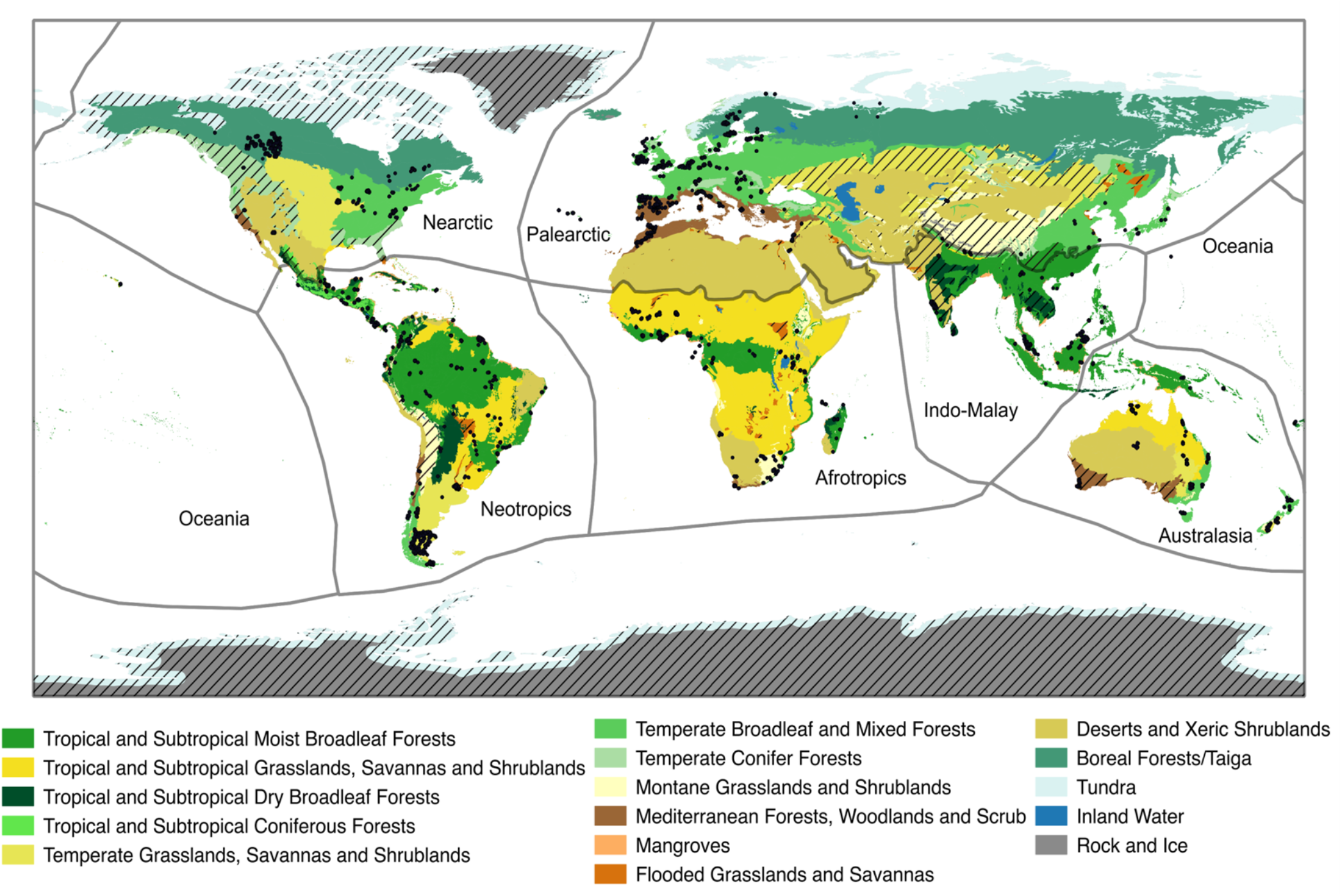
Map of regional biomes. Solid coloured areas represent biomes (see legend). Black lines separate out biogeographic realms. Black dots represent sites in the PREDICTS database used in our global model (Hudson et al 2014). Hashed coloured areas represent regional biomes that were not included in analysis due to data deficiency (see methods). Biome and biogeographic realm spatial data from (Olson *et al*., 2001).

Using the predominant land use and use-intensity classifications from the PREDICTS database (land use: primary vegetation, mature secondary vegetation, intermediate secondary vegetation, young secondary vegetation, plantation forest, pasture, cropland and urban; use intensity: minimal, light, intense) (Hudson *et al*., 2017), we created two land-use variables – *LandUse* and *UseIntensity*. We combined the three secondary vegetation classes (mature, intermediate, and young) into one and removed data from urban land-use types due to the low quantity of samples compared to other land-types (917 sites from 53 studies). Because some combinations of regional biome and *LandUse* class have low sample sizes, we tested 5 variations of this variable where categories were aggregated to find the most parsimonious grouping that still explained variation in the model well. For example, to make the *LandUse2* variable we grouped primary vegetation and secondary vegetation as ‘natural vegetation’. Full descriptions of each land-use type and the 5 variations of *LandUse* can be found in Table S2. *UseIntensity* is a combination of predominant land-use type and use-intensity. To create this variable, we grouped ‘minimal’ and ‘light’ use-intensities into ‘minimal’ and combined with groups from *LandUse3* (defined as primary vegetation, secondary vegetation, and agriculture) to give categories primary vegetation, minimal secondary vegetation, intense secondary vegetation, minimal agriculture, intense agriculture. Primary vegetation sites were not split by use intensity as there is no difference in biodiversity metrics between primary vegetation sites with differing use-intensities (Newbold *et al*., 2015).

### Statistical Analysis

We used generalised mixed-effects models (Bolker *et al*., 2009) to test for differences in species richness and total abundance in response to land use, biome, realm and regional biome (an interaction of biome and realm). Species Richness was modelled using a Poisson distribution and total abundance was log transformed (natural logarithm) and modelled with a gaussian distribution. In all models, we included study and taxa as random intercepts to account for site-level differences, data collection methods and scale differences between response metrics of Vertebrates, Invertebrates, Plants and Fungi. We ran a series of models with species richness and total abundance against each *LandUse* variable (i.e. *LandUse1, LandUse2, or LandUse3* – see the previous section) and selected the best fitting variable using Akaike Information Criterion (AIC) (Burnham, Anderson and Huyvaert, 2011). We found that *LandUse1* (Primary vegetation, Secondary vegetation, Agriculture, Plantation Forest, Cropland) had the most support and was used for remaining analyses (Table S3).

### Global analysis of regional biomes

To see if including regional biome is a more parsimonious explanation for changes in biodiversity than realm or biome, we ran a model selection process using ΔAIC values to assess the support for each variable. The fixed effect structures tested were *LandUse, LandUse:Realm, LandUse:Biome*, and *LandUse:RegionalBiome* and species richness and total abundance were used as response variables. We tested model robustness in two ways. First, we increased the sample size threshold that would allow a regional biome to be included in the model (minimum 1, 5, 25 or 50 sites in each regional biome-land use combination) and checked if the model selection process gave the same results in each case. Second, we ran a series of 100 hold-out models using the 25-site threshold dataset where each iteration randomly removed 10% of studies to see if this would change the results of model selection.

### Biome-specific case studies

As case studies, we investigated the differences in biodiversity response to land-use change between regional biomes by selecting the three biomes which had three or more regional biomes with at least 100 sites in the PREDICTS database (Fig. S1). The biomes selected were tropical and subtropical moist broadleaf forest (Tropical Forest) (3 realms; Indo-Malay: 1781 sites; Neotropics: 2274 sites; Afrotropics: 2081 sites), temperate broadleaf and mixed forests (Temperate Forest) (4 realms; Palearctic: 3986 sites; Neotropics: 577 sites; Nearctic: 856 sites; Australasia: 336 sites) and tropical and subtropical savannahs, grasslands and shrublands (Tropical Grassland) (3 realms; Afrotropic: 1663 sites; Neotropics: 190 sites; Australasia: 502 sites). For each biome and biodiversity metric (species richness and total abundance), we ran four models that used the *LandUse* or *UseIntensity* variables and tested if including *Realm*, which in this case represents regional biomes, increases model support. We then predicted the average change in species richness and total abundance, relative to primary vegetation, in each land-use type in each realm using the StatisticalModels R package (Newbold, 2015).

In addition to the biome case studies, we did a further analysis to investigate how taxonomic group further influences the responses of species richness to land-use change across regional biomes. For each biome case study, we ran one model using Species Richness as the response variable and *LandUse, Realm* and *Taxa* (Vertebrate, Invertebrate and Plant) as fixed interaction terms. As above, we predicted the average change in species richness relative to primary vegetation.

## Results

### Global analysis of regional biomes

Including regional biome as a variable improved model fit and better explained the effect of land-use change on species richness and total abundance (Fig. 2). Models with realm or biome alone had lower support and this was consistent across 100 iterations of hold-out models (Fig. 2). Increasing the sample size threshold for regional biomes did not change the overall outcome (Table S4). Furthermore, marginal R^2^ values showed that regional biome explained more variation in both species richness and total abundance than biome or realm interacting with land use (Table S4).

**Figure 2.**
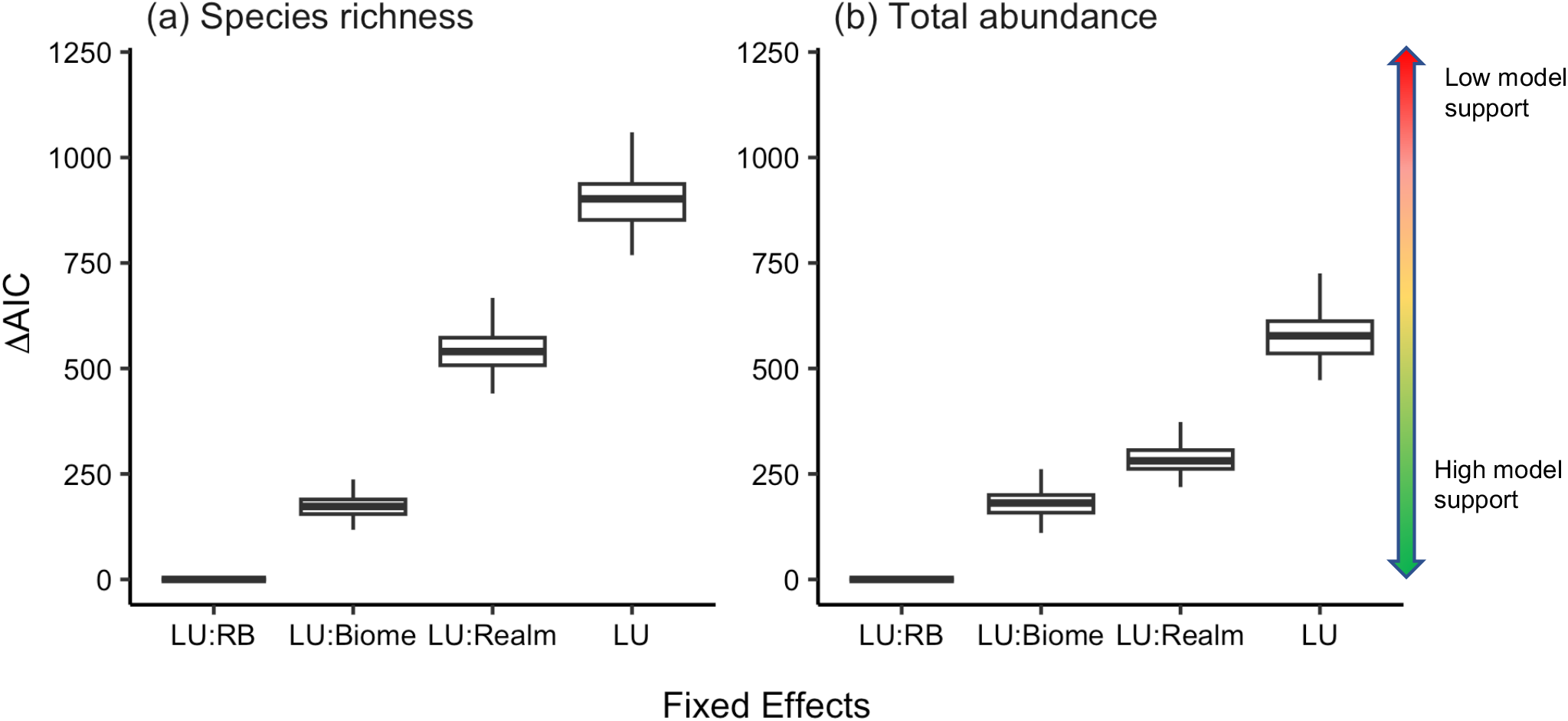
Change in Akaike’s Information Criterion (ΔAIC) for Generalised Linear Mixed Models (GLMMs) predicting change in a global dataset of (a) species richness and (b) total abundance consistently suggests models including regional biome have highest support when modelling response to land-use change. Four GLMMs with differing fixed-effect structures containing Land Use (LU) interacting with Regional Biome (RB), Biome or Realm were compared using AIC values. A model with low AIC is considered to have higher support. ΔAIC is calculated as the difference in AIC from the lowest scoring model. The box and whisker plots represent a summary of ΔAIC values of 100 hold-out models, where each iteration removed 10% of studies at random as a test of model robustness. Importantly, for a given dataset in this hold-out analysis, LU:RB model had the lowest AIC in 100% of iterations. The PREDICTS dataset was subset to only include data from regional biomes with 25 data points per combination of regional biome and land-use type. Land Use is a discrete variable with Primary Vegetation, Secondary Vegetation, Plantation Forest, Pasture and Cropland as its categories (see Table S2 for descriptions).

### Biome-specific case studies

We found that species richness and total abundance in each regional biome of tropical forest showed a distinct response to land-use change. Species richness reduced in human-dominated land-use types compared to primary vegetation in all realms but to different degrees at each land-use type (Fig. 3a). The largest loss of species richness was seen in Afrotropical cropland (-53% reduction compared to primary vegetation), followed by Indo-Malayan cropland (-50%) and plantation forest (-43%). Conversely, the Neotropics did not show a significant species richness reduction in cropland and a barely significant reduction in plantation forest (-20%), with species richness for this regional biome being worst affected in pasture land-use types (-30%). When predicting responses of total abundance to land-use change in the tropical forest biome there was much higher uncertainty. The negative responses to land-use type seen in the species richness models did not always carry over to the abundance models, for example there was no sign of a negative response in Neotropical pasture. However, total abundance was significantly lower in Indo-Malay plantation forest compared to other regional biomes (Fig. S2a). For the models including *UseIntensity*, a negative response to increased disturbance can be observed in Indo-Malayan and Afrotropical regional biomes, with declines >50% compared to primary vegetation in both species richness and total abundance (Fig. 3b, Fig. S2b). This extreme response is not seen in the Neotropical regional biome.

**Figure 3.**
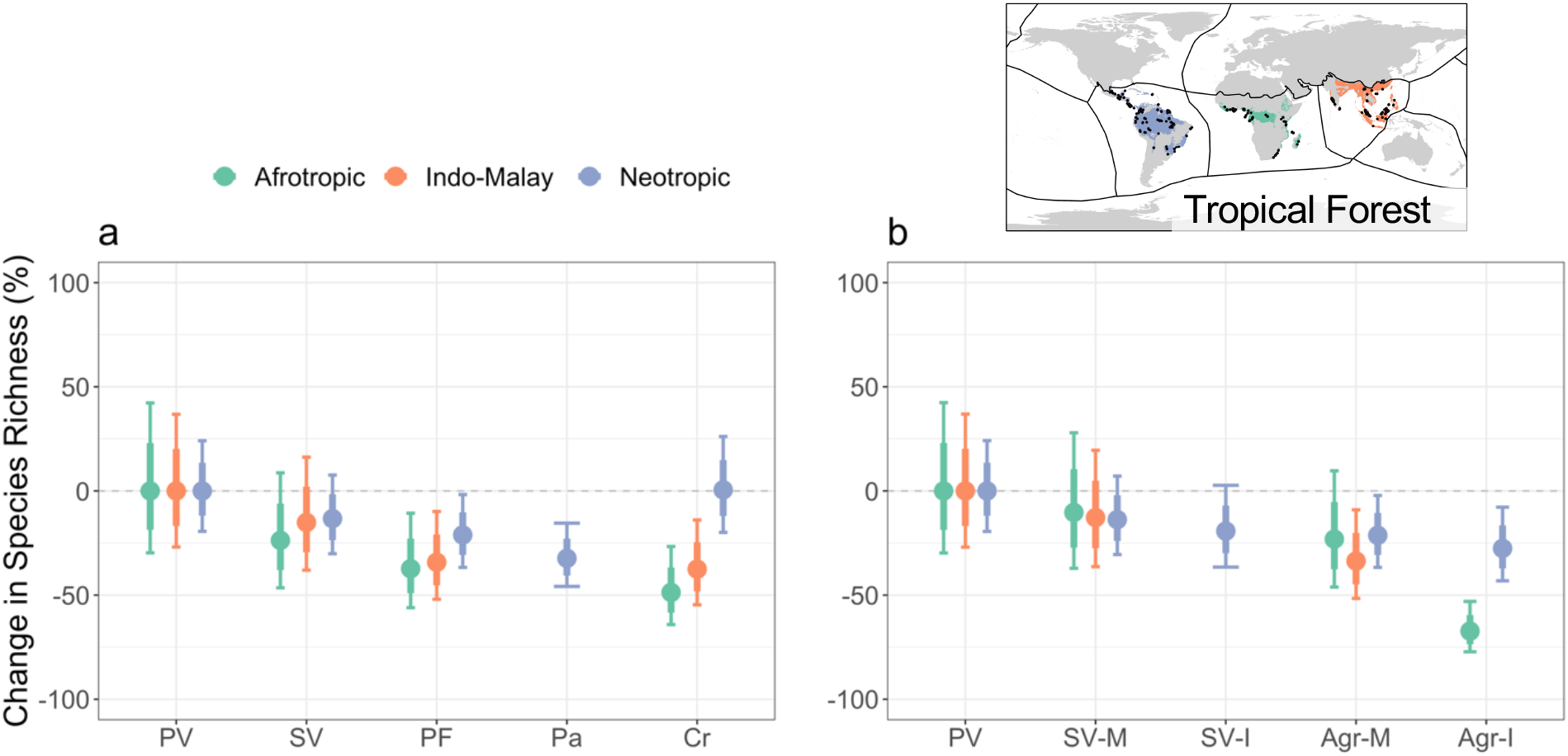
Responses of biodiversity to land-use change are unique across tropical forest regional biomes. The results of GLMMs predicting the response of species richness across land-use types and land-use intensity, when compared to primary vegetation (PV) across tropical forest regional biomes. Responses of species richness of plants and animals were predicted across land-use types (a) including secondary vegetation (SV), plantation forest (PF), pasture (Pa) and cropland (Cr). Responses were also predicted for two levels of land-use intensity: minimal (M) or intense (I) in secondary vegetation (SV) and agriculture (Agr) land use types (b). Each point represents mean prediction, with 75% confidence intervals (thick whiskers) and 95% confidence intervals (thin whiskers). Responses are considered significantly different if the 75% confidence intervals do not overlap.

In temperate forest regional biomes, species richness and total abundance both showed low to no response in human-dominated land-use types compared to primary vegetation (Fig. 4a, Fig. S2c). The exception is the Palearctic regional biome, where total abundance reduced by 55% in cropland land-use types. There were also reductions in species richness in plantation forest and cropland in this regional biome, but the upper confidence limits just surpassing 0 (Fig. 4a). This was the only response from any temperate forest regional biome that showed a significant change from primary vegetation (95% confidence intervals do not include 0). Furthermore, predictions for change in species richness and total abundance have high levels of uncertainty for all realms except palearctic.

**Figure 4.**
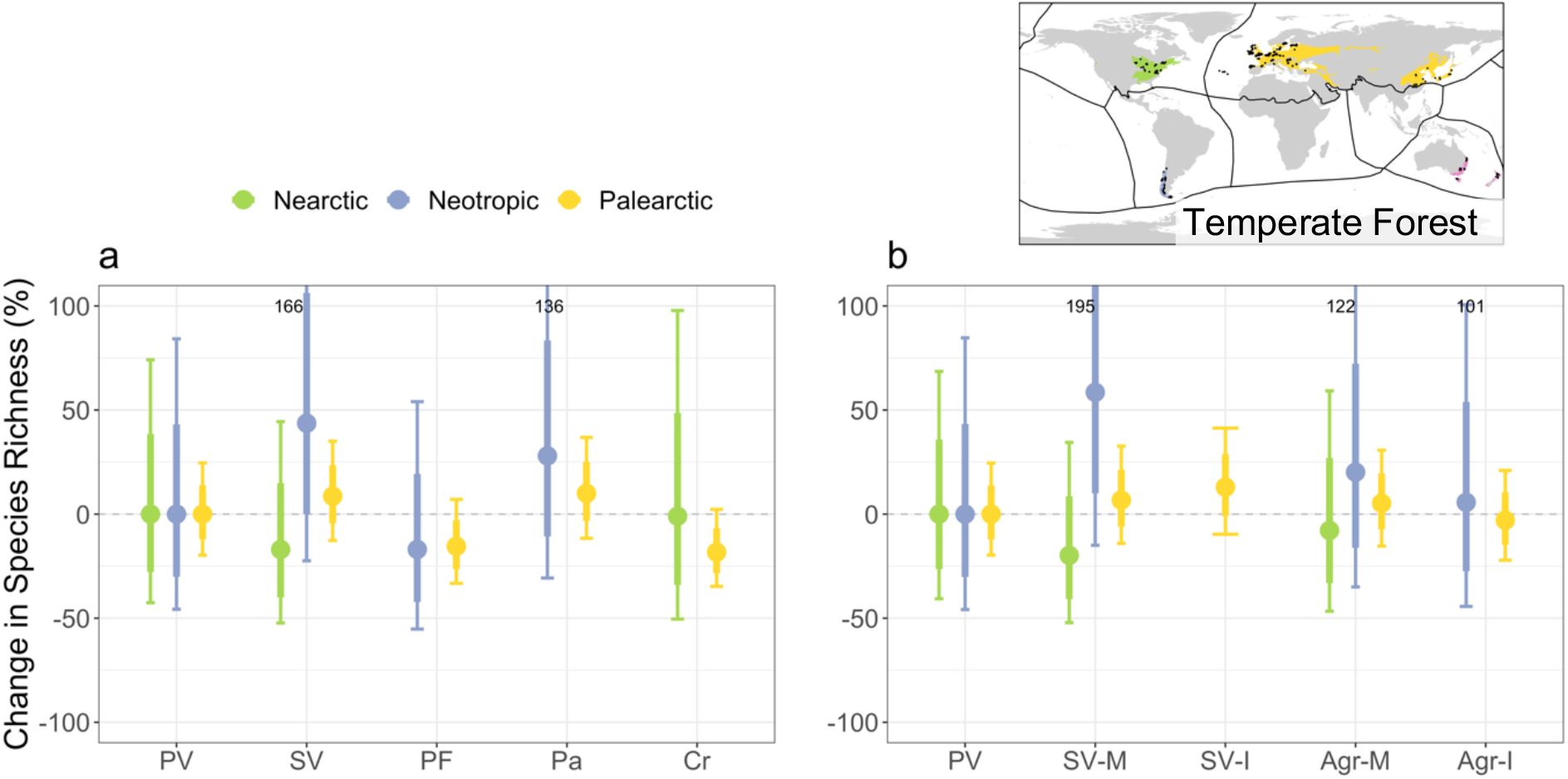
Responses of biodiversity to land-use change are hard to predict unique across temperate forest regional biomes. The results of GLMMs predicting the response of species richness across land-use types and land-use intensity, when compared to primary vegetation (PV) across temperate forest regional biomes. Responses of species richness of plants and animals were predicted across land-use types (a) including secondary vegetation (SV), plantation forest (PF), pasture (Pa) and cropland (Cr). Responses were also predicted for two levels of land-use intensity: minimal (M) or intense (I) in secondary vegetation (SV) and agriculture (Agr) land use types (b). Each point represents mean prediction, with 75% confidence intervals (thick whiskers) and 95% confidence intervals (thin whiskers). To keep scales consistent, upper 95% CIs greater than 100 have been included as text instead. Responses are considered significantly different if the 75% confidence intervals do not overlap.

Similar to temperate forest, biodiversity metrics in most tropical grassland regional biomes showed little or no response to any land-use type when compared to primary vegetation (Fig. 5). Predictions of species richness and total abundance change showed a high degree of uncertainty across all regional biomes, except for the Afrotropics, where there was a strong response to intense agriculture (-67%), although this was the only regional biome with enough datapoints to predict in this category so cannot be compared (Fig. 5b, Fig. S2f). Species richness in Australasia showed negative responses to pasture and minimally-used agricultural land, but with the upper confidence interval just passing zero in both cases (Fig. 5a, b).

**Figure 5.**
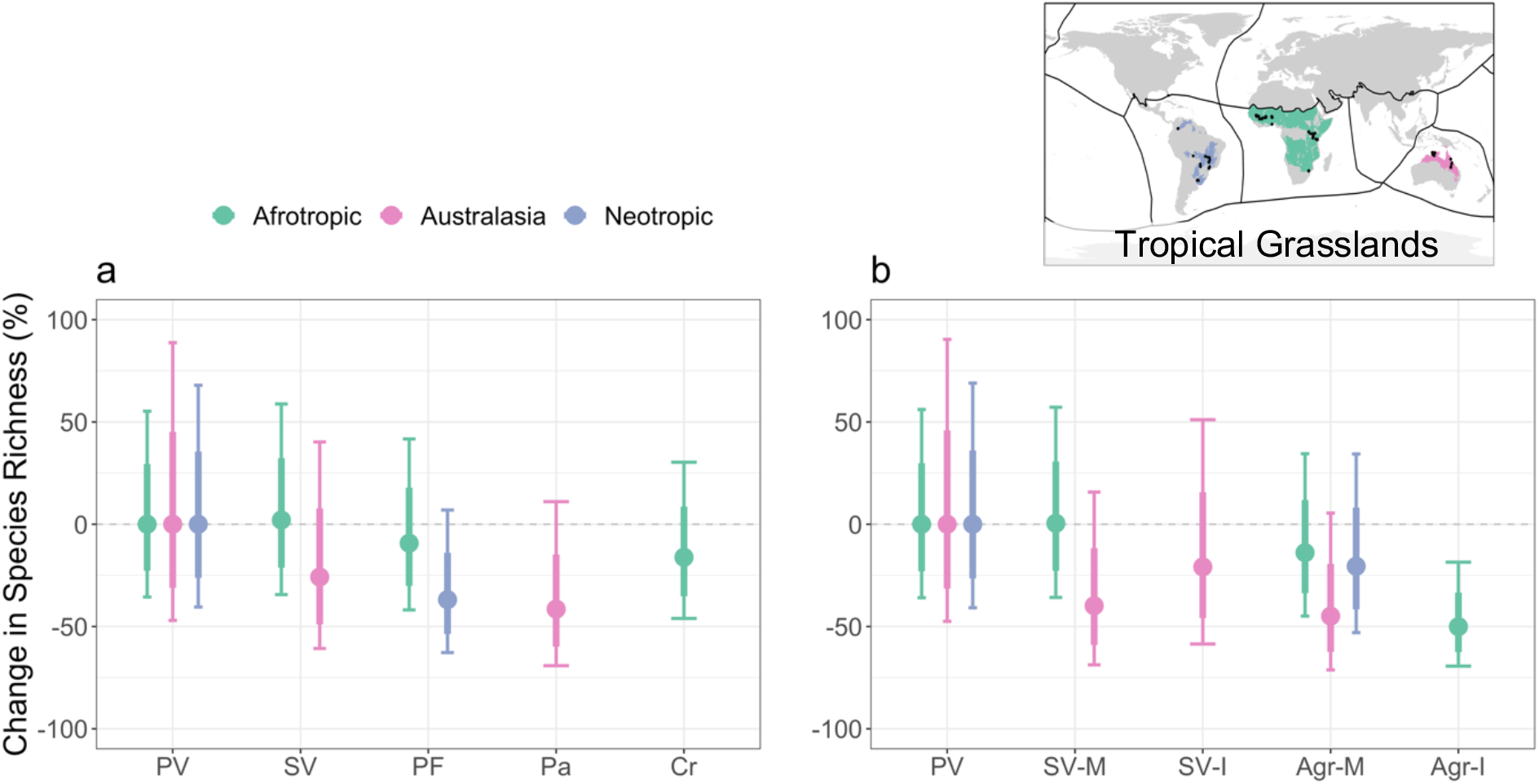
Responses of biodiversity to land-use change are hard to predict unique across tropical grassland regional biomes. The results of GLMMs predicting the response of species richness across land-use types and land-use intensity, when compared to primary vegetation (PV) across tropical grassland regional biomes. Responses of species richness of plants and animals were predicted across land-use types (a) including secondary vegetation (SV), plantation forest (PF), pasture (Pa) and cropland (Cr). Responses were also predicted for two levels of land-use intensity: minimal (M) or intense (I) in secondary vegetation (SV) and agriculture (Agr) land use types (b). Each point represents mean prediction, with 75% confidence intervals (thick whiskers) and 95% confidence intervals (thin whiskers). To keep scales consistent, upper 95% CIs greater than 100 have been included as text instead. Responses are considered significantly different if the 75% confidence intervals do not overlap.

Including taxon as an interaction term reveals distinct responses to land-use change across different taxonomic groups within regional biomes (Fig. 6). In tropical forest, there is still a general trend of strong negative responses to human-dominated land-use types, but for some regional biomes there is high variation in responses. For example, afrotropical vertebrates have no observable response to cropland and plantation forest, but plants in the same regional biome display strong reductions in species richness as these land-use types. Conversely, in Neotropical tropical forest, Vertebrates and Invertebrates show a negative response to pasture land, but plants do not. In temperate forest, the addition of taxon groups highlights the level of incomplete sampling in the dataset but reveals strong responses of vertebrates and plants to land-use change in the Palearctic regional biome that are missed in the unified model (as seen in Fig. 4). In tropical grassland, we observe the opposite effect, where there are no strong responses predicted to any land-use type and increased uncertainty in predictions, due to much lower sampling effort.

**Figure 6.**
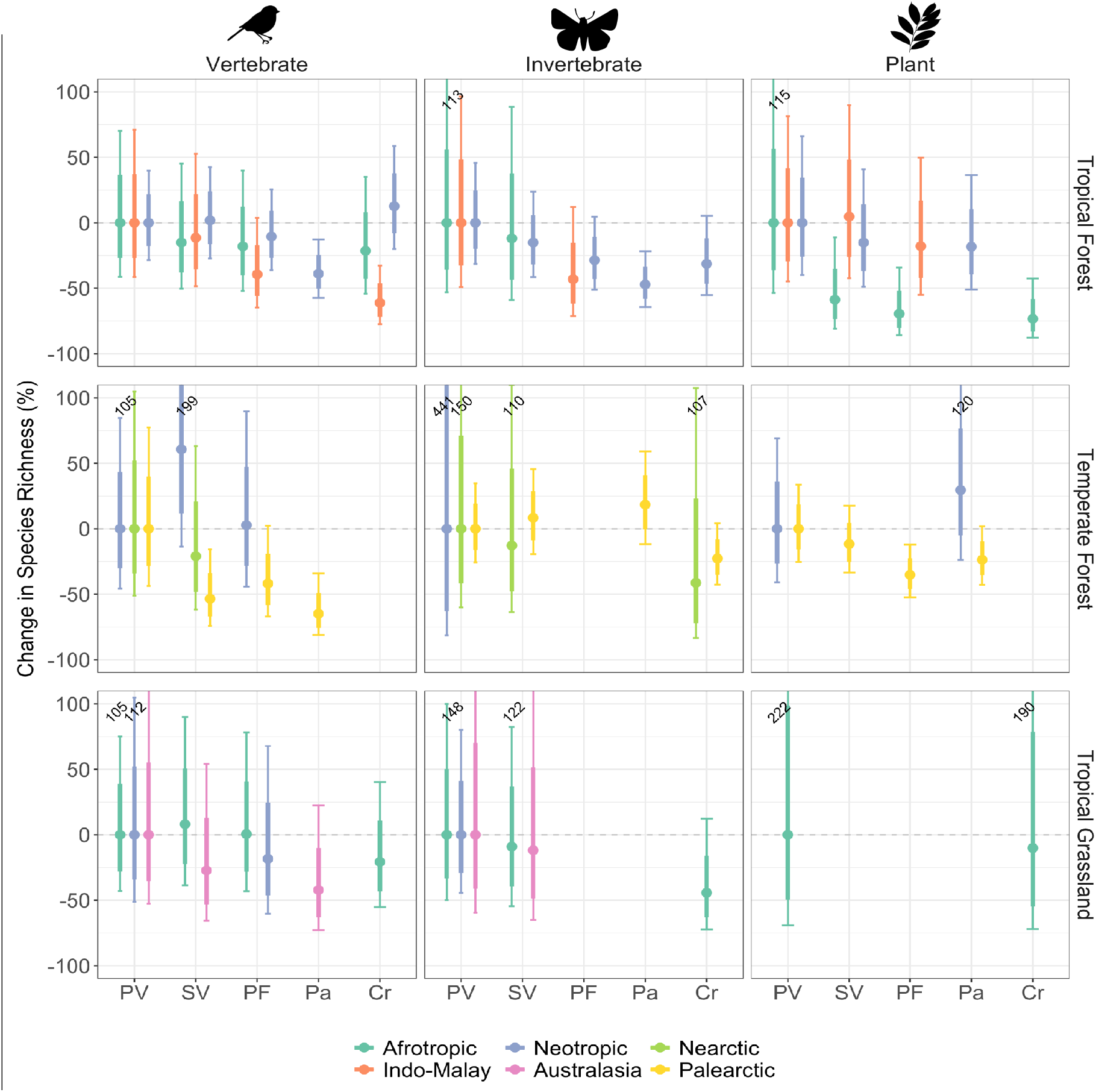
Responses of species richness in individual taxa groups to land-use change across regional biomes. The results of 6 GLMMs predicting the response of species richness across land-use types and realm across three biomes (Tropical Forest, Temperate Forest and Tropical grassland) and three taxon groups (Vertebrate, Invertebrate and Plants) when compared to primary vegetation (PV) across tropical grassland regional biomes. Responses of species richness were predicted across disturbed land-use types including secondary vegetation (SV), plantation forest (PF), pasture (Pa) and cropland (Cr). Each point represents mean prediction, with 75% confidence intervals (thick whiskers) and 95% confidence intervals (thin whiskers). To keep scales consistent, upper 95% CIs greater than 100 have been included as text instead. Responses are considered significantly different if the 75% confidence intervals do not overlap.

## Discussion

### Global analysis of regional biomes

Our results show that including regional biome when modelling the global effects of land use on biodiversity can increase explanatory power. There is a need for a biodiversity monitoring spatial framework that finds the balance between having feasible data collection requirements and adequate global coverage, at a realistic scale that captures true variation in ecological resilience to disturbance (Ingram *et al*., 2021). It has been previously shown that biodiversity shows different responses to disturbance across biomes (Greenville *et al*., 2018; Blowes *et al*., 2019; Newbold *et al*., 2020) and across realms (Gibson *et al*., 2011; Phillips, Newbold and Purvis, 2017; Lambertini, 2020), but for the first time, we have shown the important interaction between biome and realm. The combination of these two spatial delineations gives us more information about ecological processes. This result is important for biodiversity monitoring and conservation interventions as not only are the magnitude of biodiversity responses to anthropogenic pressure different between regions but that threatening processes (i.e., land-use types) are affecting regional biomes differently.

### Biome-specific case studies

Whilst our global models suggest that regional biome can modulate the effect of land-use change on species richness and abundance on a global scale, a more focused analysis found that the influence of regional biome is stronger in some biomes than others. Generally, species richness and total abundance in tropical biomes showed negative responses to disturbed land-use types, the degree of which changed with realm and taxonomic group. In temperate forest, however, there was no significant response to land-use change overall, excluding the palearctic realm, particularly vertebrates. The observed difference between tropical and temperate regional biomes is expected; responses to land-use change are stronger in tropical biomes compared to temperate biomes, especially tropical forest (Newbold *et al*., 2020). However, ecoregions in tropical biomes are considered more ecologically distinct than neighbouring ecoregions compared to those in temperate biomes (Smith *et al*., 2018). As each regional biome is a group of many ecoregions, it would follow that predicted responses to disturbance in tropical regional biomes would contain more uncertainty than temperate, but the opposite is true for our case studies. The high uncertainty in predicted responses produced by the temperate forest and tropical grassland models can be partly explained by low sample sizes and the high variation in responses of taxonomic groups. Predictions in palearctic temperate forest have very low uncertainty compared to all other regional biomes and is also the most extensively sampled regional biome in the PREDICTS database (Table S1).

For tropical forest regional biomes, our model predictions suggest Indo-Malayan biodiversity responds most strongly to disturbance, followed by the Afrotropics. There was no significant change in species richness in the Neotropics tropical forest in any land-use type except for pasture. Adding taxon to the model showed that these responses are nuanced and driven by different taxonomic groups, but the patterns remain largely the same. Biodiversity in Asian tropical forest is repeatedly found to be extremely sensitive to land-use change and disturbance (Sodhi *et al*., 2009; Gibson *et al*., 2011; Phillips, Newbold and Purvis, 2017). The islands of south east Asia have a rich geographical history which has led to this region having the highest number of endemic species and biodiversity hotspots (Myers *et al*., 2000; Sodhi *et al*., 2004) compared to other tropical regions. This area also has the highest proportion of species under threat in the world (Allan *et al*., 2019). This high baseline for species richness combined with high levels of specialisation may partly explain why the response to land-use change is so pronounced in this regional biome. Response metrics in Neotropical tropical forest have the smallest response to disturbed land-use types compared to similar regional biomes. However, temporal trends from the Living Planet Index suggest that populations of vertebrates in south America have decreased by 94% since 1970 (Lambertini, 2020). In addition, other studies in this region have shown a decline in biodiversity due to land-use change (Bogoni, Peres and Ferraz, 2020; Gaona *et al*., 2021; Quintero *et al*., 2023). The difference is a demonstration of the importance of testing multiple data types and of assessing the regional biome framework using time-series biodiversity data and across multiple threat gradients in the future. Further research is needed to understand how biological and taxonomic differences between these tropical forest realms might be impacting sensitivity to land-use change.

The intensity and history of habitat disturbance in each regional biome will also impact the observed response of biodiversity and may even cause biases in our data set. There is a strong focus on palm oil as a driver of biodiversity loss in Asian tropics, which may lead to a publication bias in this area. Temperate forest biomes have a long history of human habitation and extinction events, meaning the baseline in primary vegetation is much lower in these regional biomes compared to less historically disturbed biomes (Monsarrat and Svenning, 2022). The lack of variation in responses observed in temperate forest regional biomes may well be because of this lower baseline in primary vegetation. Indeed, trends of vertebrate abundance have been increasing in the northern hemisphere since 1970 (Leung, Greenberg and Green, 2017) as well as forest cover (Song *et al*., 2018). The drivers that cause differences in sensitivity to habitat disturbance are related to both biogeographic and human history, splitting the world into regional biomes can further account for these differences.

Although regional biomes may be a valuable spatial monitoring unit, our models were limited by sample size for some regional biomes. The PREDICTS database is the most extensive collection of terrestrial biodiversity records available, but there are still gaps. Under-sampling was pronounced in non-forested temperate biomes like Montane Grasslands and Shrublands, and realms were unevenly sampled, for example studies from Palearctic temperate forest dominate the database but there are no studies at all on Palearctic temperate grasslands (Table S1). Furthermore, taxonomic under sampling was highlighted in our model that included taxonomic group. For example, Fig. 3 suggests that afrotropical tropical forests have a greater response to Cropland compared to the neotropical regional biome. However, this response is driven by Plants in the Afrotropics and Invertebrates and Vertebrates in the Neotropics, making the impact of cropland hard to compare. Prioritising data collection in under-represented regional biomes and taxonomic groups highlighted here would enhance progress towards an effective monitoring framework (Ingram *et al*., 2021). Furthermore, this study focuses only on terrestrial regional biomes, as these are more extensively sampled and we were specifically testing the effect of land-use change on species richness and abundance. We acknowledge that marine biodiversity is showing stronger responses to disturbance than terrestrial (Blowes *et al*., 2019), and suggest a similar monitoring framework could be adapted for marine ecosystems.

## Conclusion

Our results show that the taxa-regional biome unit has the potential to be a powerful framework for monitoring biodiversity and implementing conservation action. Resources for data collection and biodiversity monitoring on the ground are sparse, and the regional biome framework implies that monitoring studies could be more evenly spread across regional biomes and taxonomic groups, optimising resource allocation for data collection. Here we have highlighted that there is within-biome variation in how biodiversity is responding to disturbance. We show that the difference between realms may be more important in tropical biomes than temperate, and that spatial resolution should be carefully considered in any attempts at monitoring global biodiversity trends. Although more data are needed, regional biomes have the potential to be a meaningful scale at which to prioritise monitoring. These results will be beneficial for informing upcoming policy discussions such as the CBD’s COP16 on the best way to monitor progress towards biodiversity targets and effect change on the ground.

## Data Accessibility Statement

All data used in this manuscript is downloaded from open-source locations. The PREDICTS database can be downloaded from the Natural History Museum Data portal (https://data.nhm.ac.uk/dataset/the-2016-release-of-the-predicts-database).

Geographical data used to delineate regional biome boundaries in this manuscript can be downloaded from The Nature Conservancy’s Geospatial Conservation Atlas (https://geospatial.tnc.org/datasets/b1636d640ede4d6ca8f5e369f2dc368b/about).

All R code used to run analysis presented in this manuscript is accessible in a GitHub repository (not included here to maintain anonymity)

