## SupplementaryInformation for "Biodiversity shows unique responses to land-use change across regional biomes"

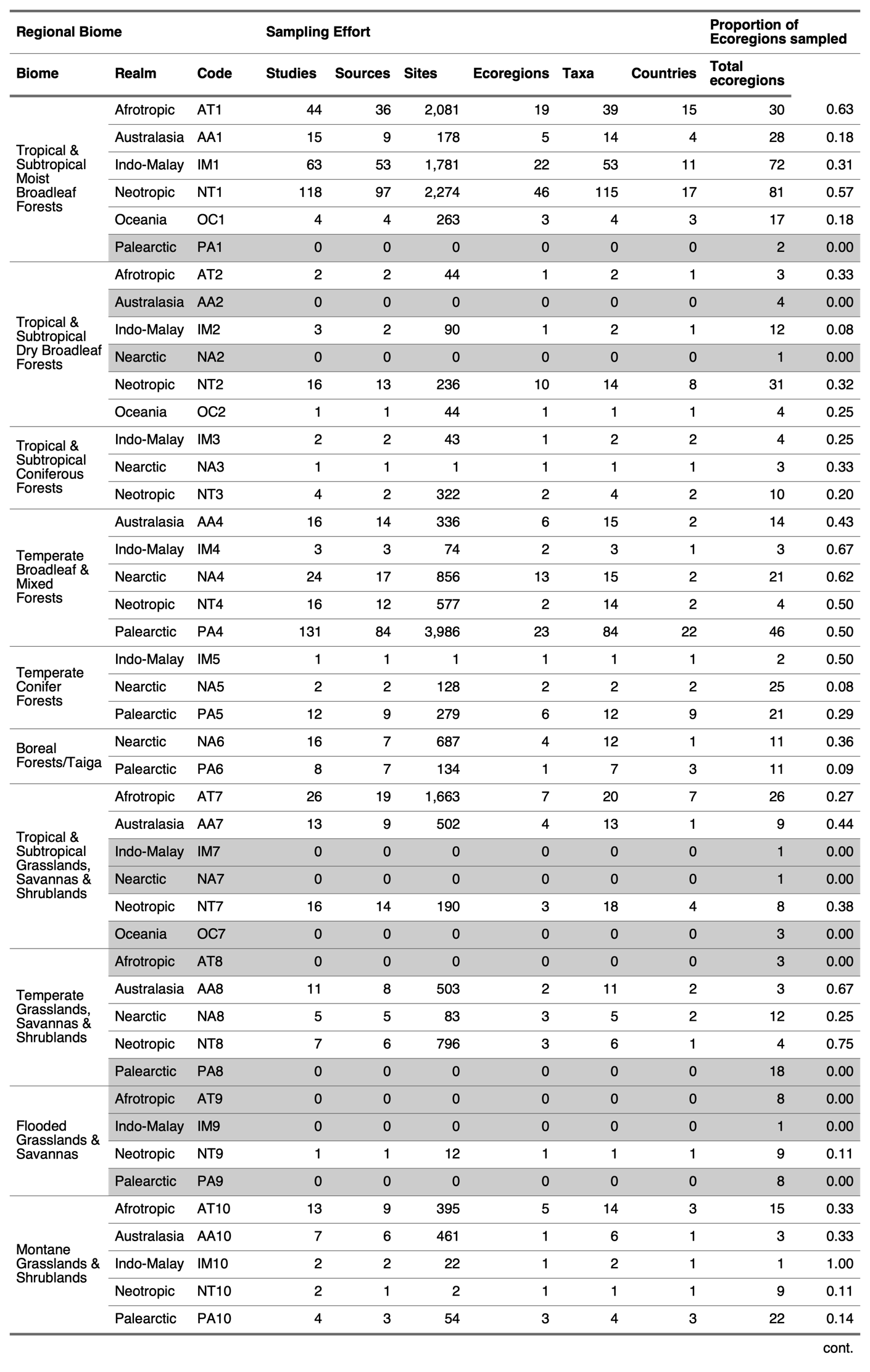

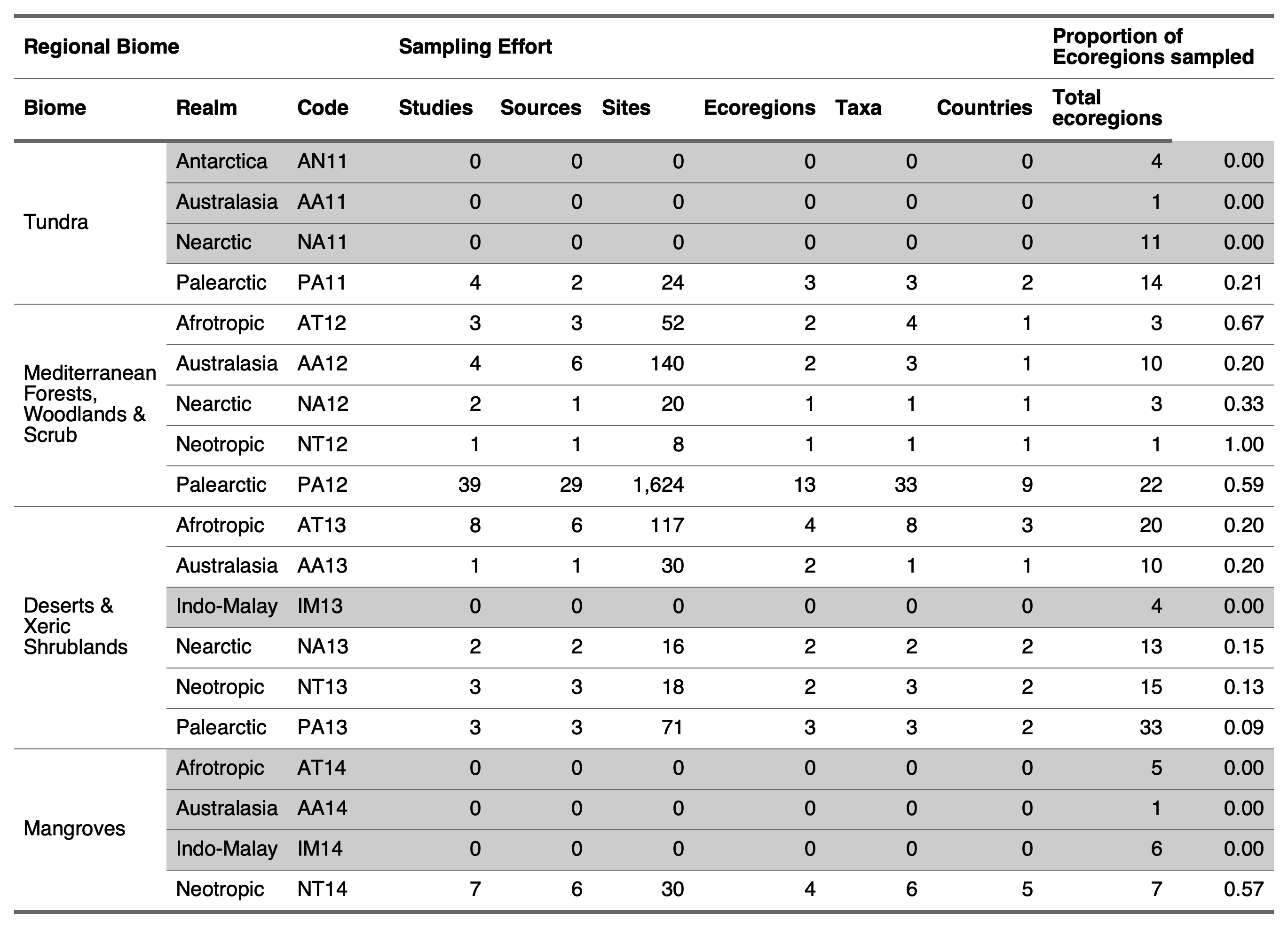
**Table S1**. Table of regional biomes and their representation in the PREDICTS database. Studies, sources and sites show the number of sources and sample sites for each regional biome in the PREDICTS database whilst Ecoregions, Taxa and Countries columns represent the number of ecoregions, taxonomic groups and countries covered. The final columns show the actual number of ecoregions in each regional biome (calculated from Olson et al., 2001), and the proportion of ecoregions covered by the PREDICTS database. Grey rows represent regional biomes that are unsampled by the PREDICTS database.

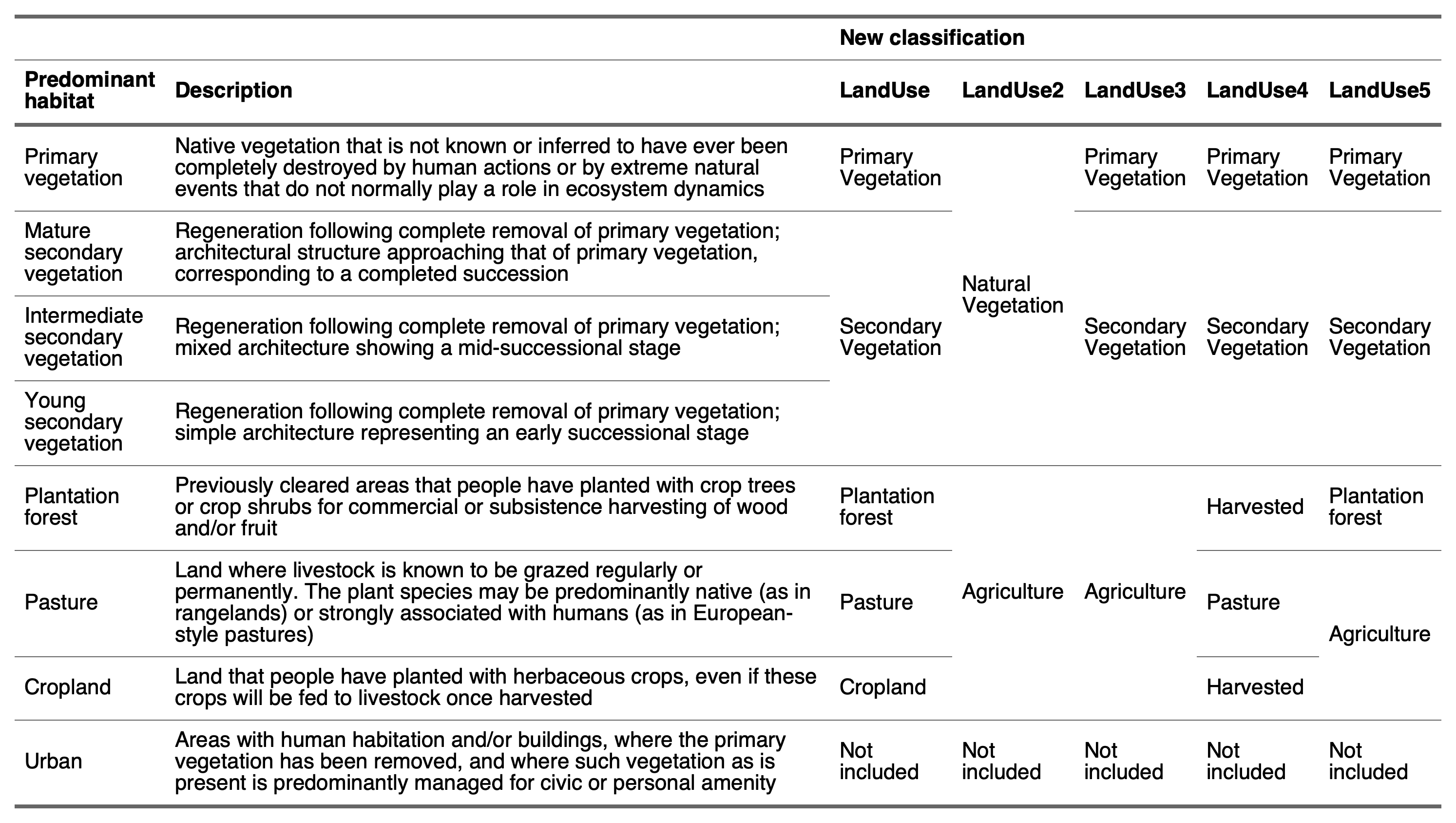

**Table S2.** Description of land-use types in the PREDICTS database and different grouping methods tested in model selection. See Hudson *et al.* (2014) supplementary information for full details of land use classification.

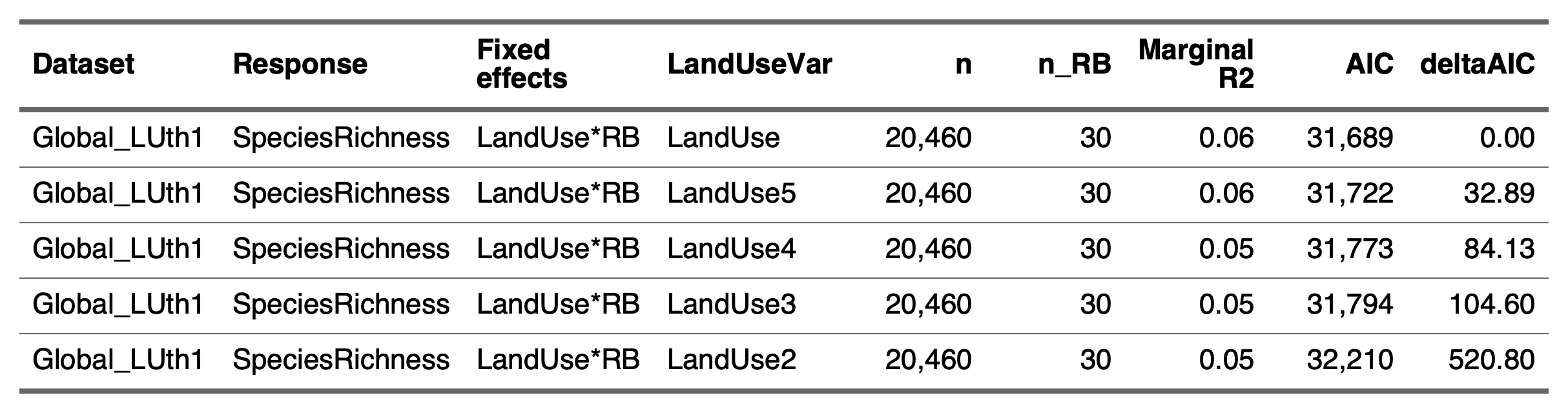

**Table S3.** *LandUse* variable selection for global model. A model was run on the same dataset using each of the different *LandUse* variables, described in Table S2. Model selection criteria include Marginal R2, a measure of variance explained by the fixed effects, and change in AIC (deltaAIC).

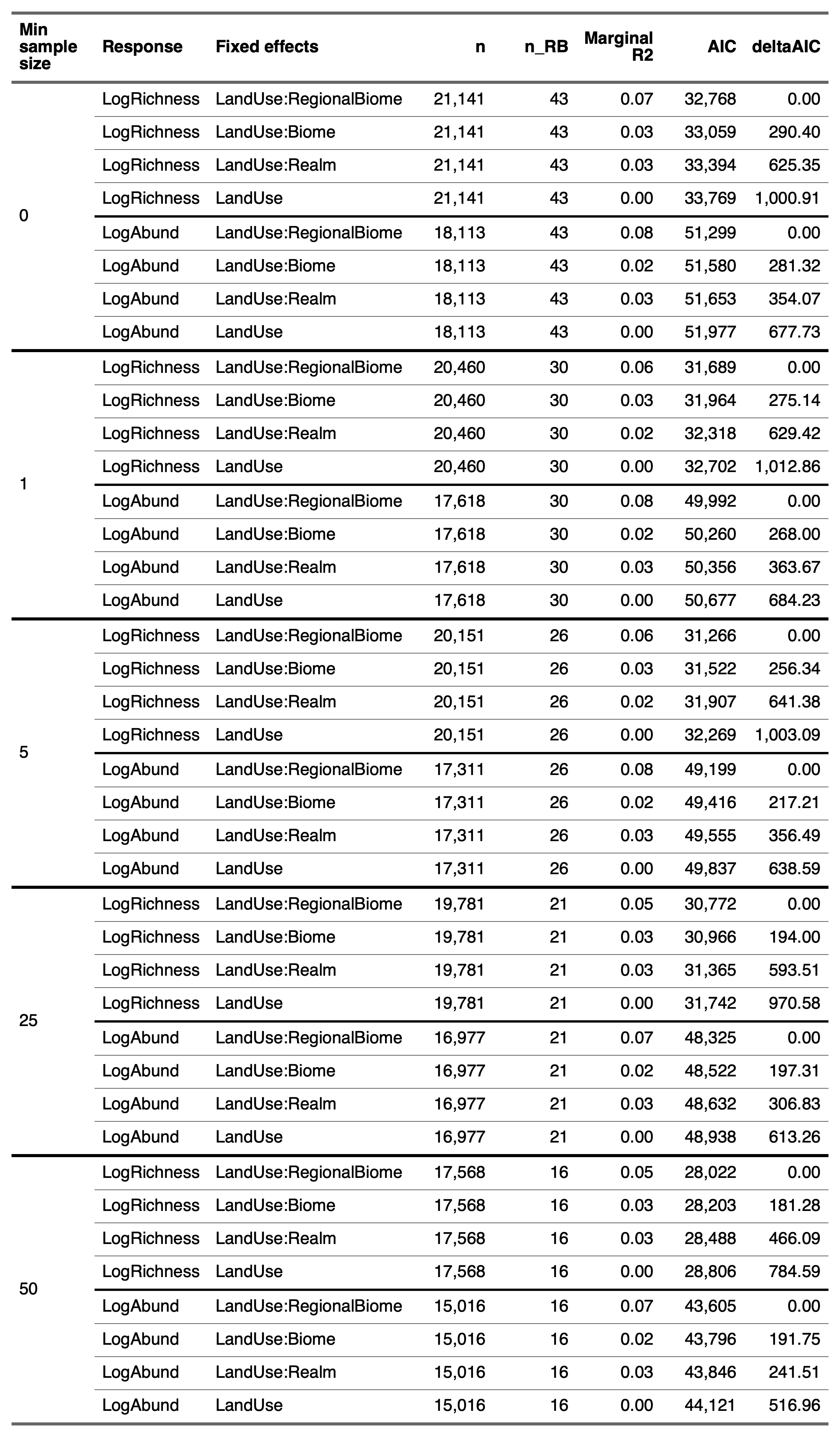

**Table S4**. Impact of changing minimum sample size needed to exclude a regional biome from the global model. Regional biomes were considered data deficient if the number of samples in either primary vegetation or any of the human dominated land-use types did not meet a certain threshold. The impact of changing this threshold was considered by running the same set of models on data subsets created from 5 different threshold levels (0,1,5,25,50), and asking if this would change the highest supported model, based on change in AIC values and marginal R2 values. In this case, the highest supported model was consistent across all threshold values.

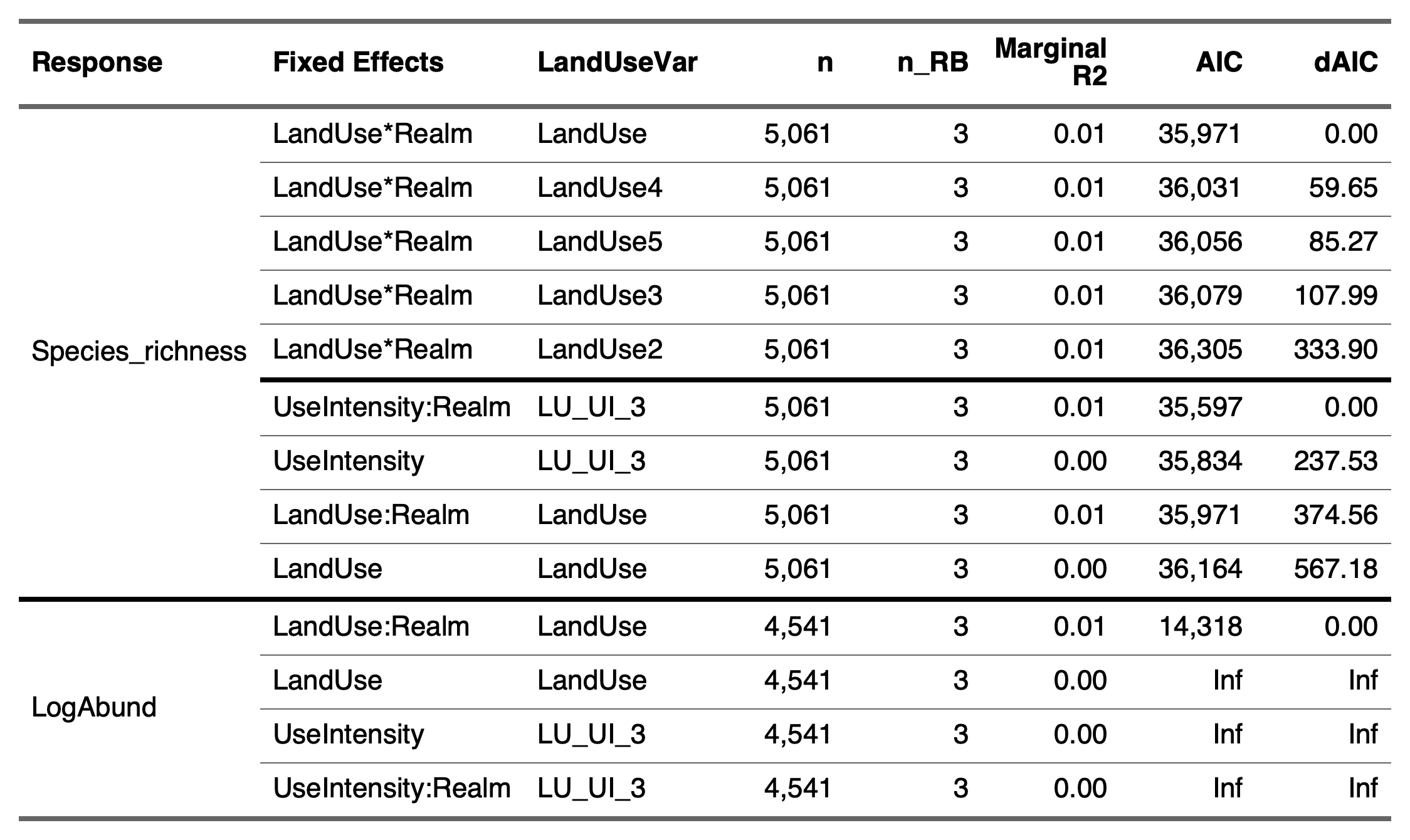

**Table S5**. Model selection results for Tropical & Subtropical Moist Broadleaf forest. For descriptions of Land Use fixed effects see Table S2. Models with AIC value of ‘Inf’ indicate models that failed to converge.

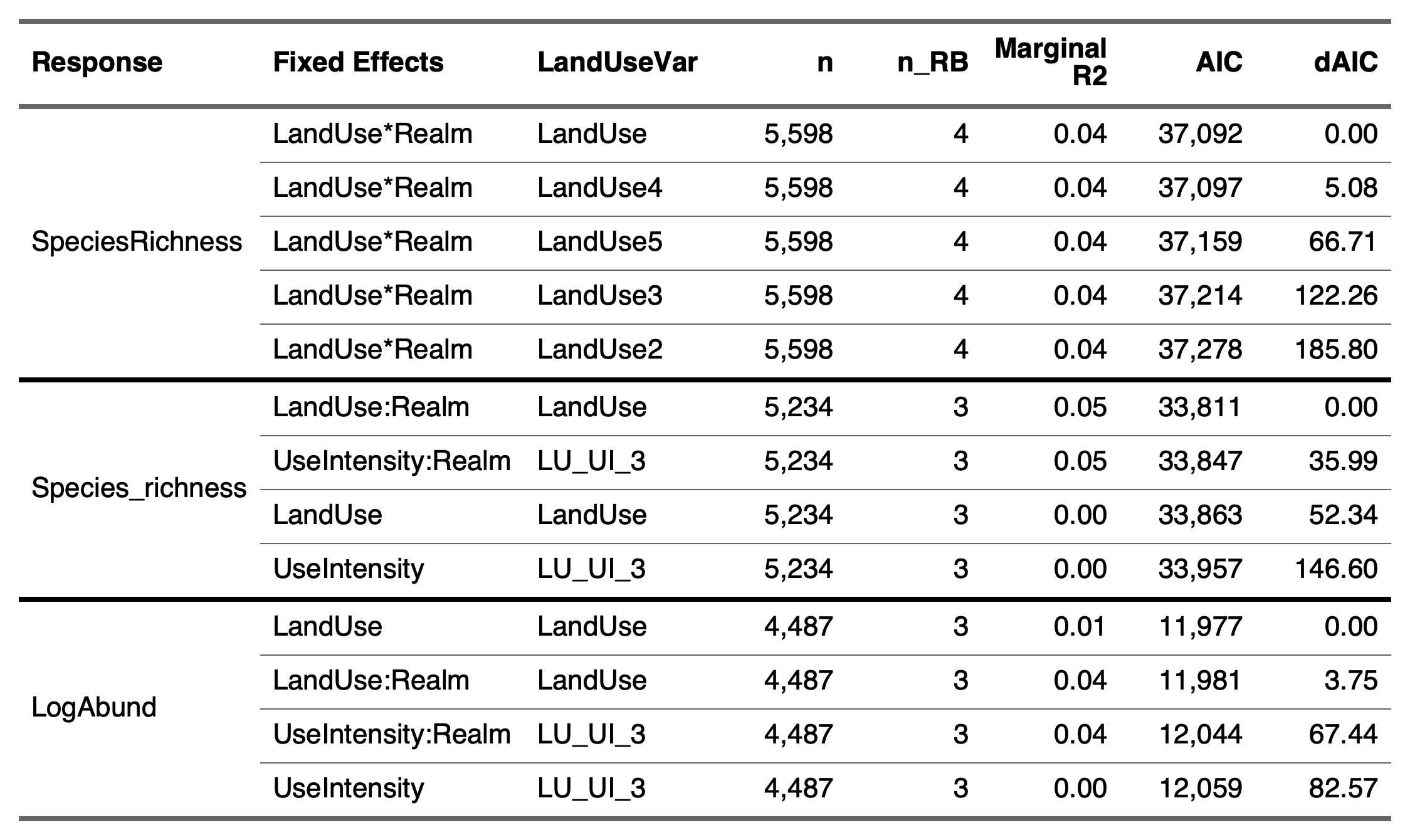

**Table S6.** Model selection results for Temperate Broadleaf and Mixed Forests. For descriptions of Land Use fixed effects see Table S2.

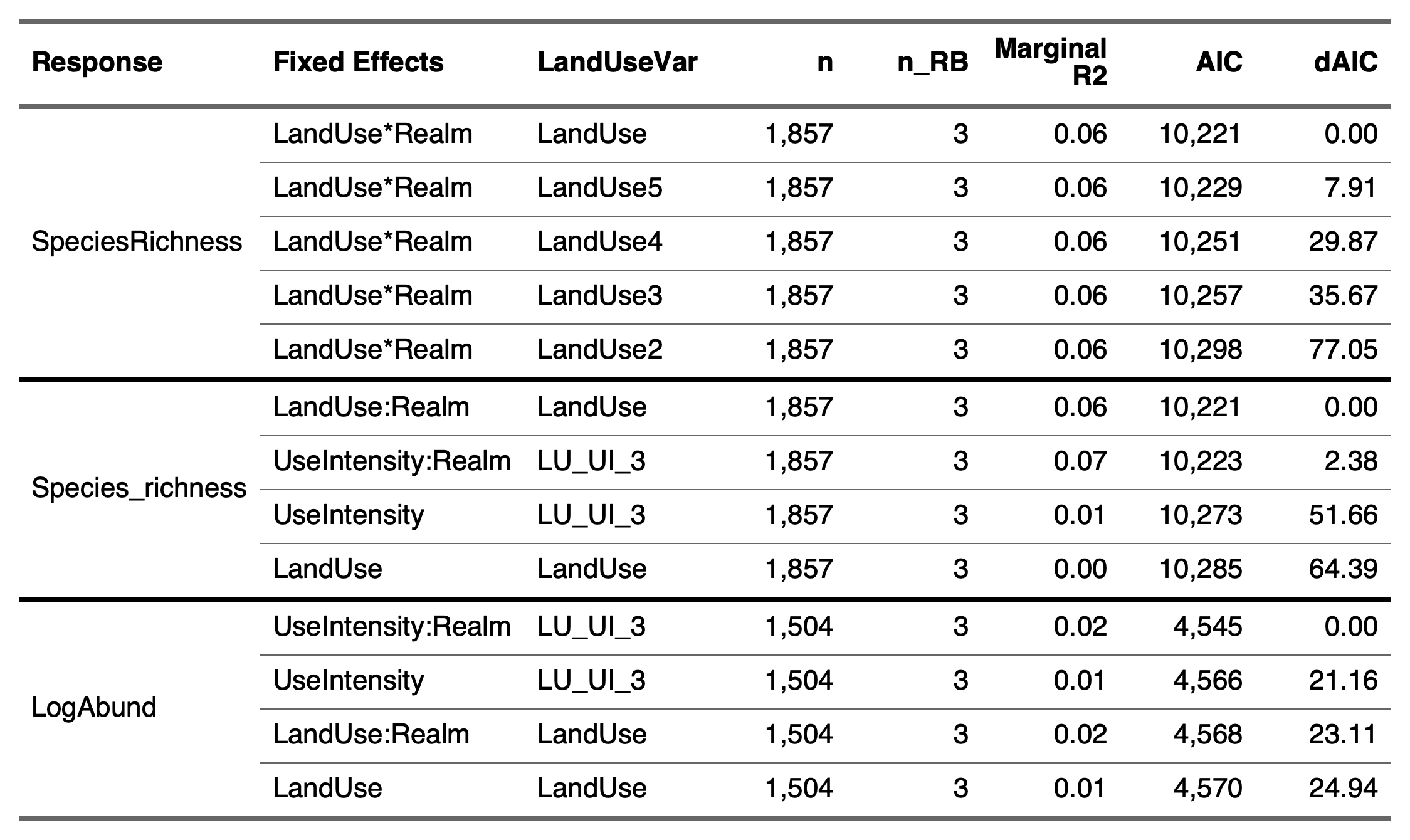

**Table S7.** Model selection results for Tropical & Subtropical Grasslands, Savannas & Shrublands. For descriptions of Land Use fixed effects see Table S2.

**Figure S1.** **Three case-studies of regional biomes: distribution of data.** The PREDICTS database was subset into studies from three biomes that included at least 25 data points in each land use category from a minimum of three biogeographic realms. The figures show ranges of regional biomes, coloured by realm, for (a) Tropical Forest, (b) Temperate Forest and (c) Tropical grassland. Black dots represent a data point (record of species richness or total abundance) from the PREDICTS dataset. Regional biome distribution data from (Olson *et al.*, 2001).

**Figure S2.** **Responses of total abundance to land-use change across regional biomes.** The results of GLMMs predicting the response of total abundance across land-use types and land-use intensity, when compared to primary vegetation (PV) across tropical forest (top row), temperate forest (middle row) and tropical grassland (bottom row) regional biomes. Responses of total abundance of plants and animals were predicted across land-use types including secondary vegetation (SV), plantation forest (PF), pasture (Pa) and cropland (Cr) (a,c,e). Responses were also predicted for two levels of land-use intensity; minimal (M) or intense (I) in secondary vegetation (SV) and agriculture (Agr) land use types (b,d,f). Each point represents mean prediction, with 75% confidence intervals (thick whiskers) and 95% confidence intervals (thin whiskers). Responses are considered significantly different if the 75% confidence intervals do not overlap.

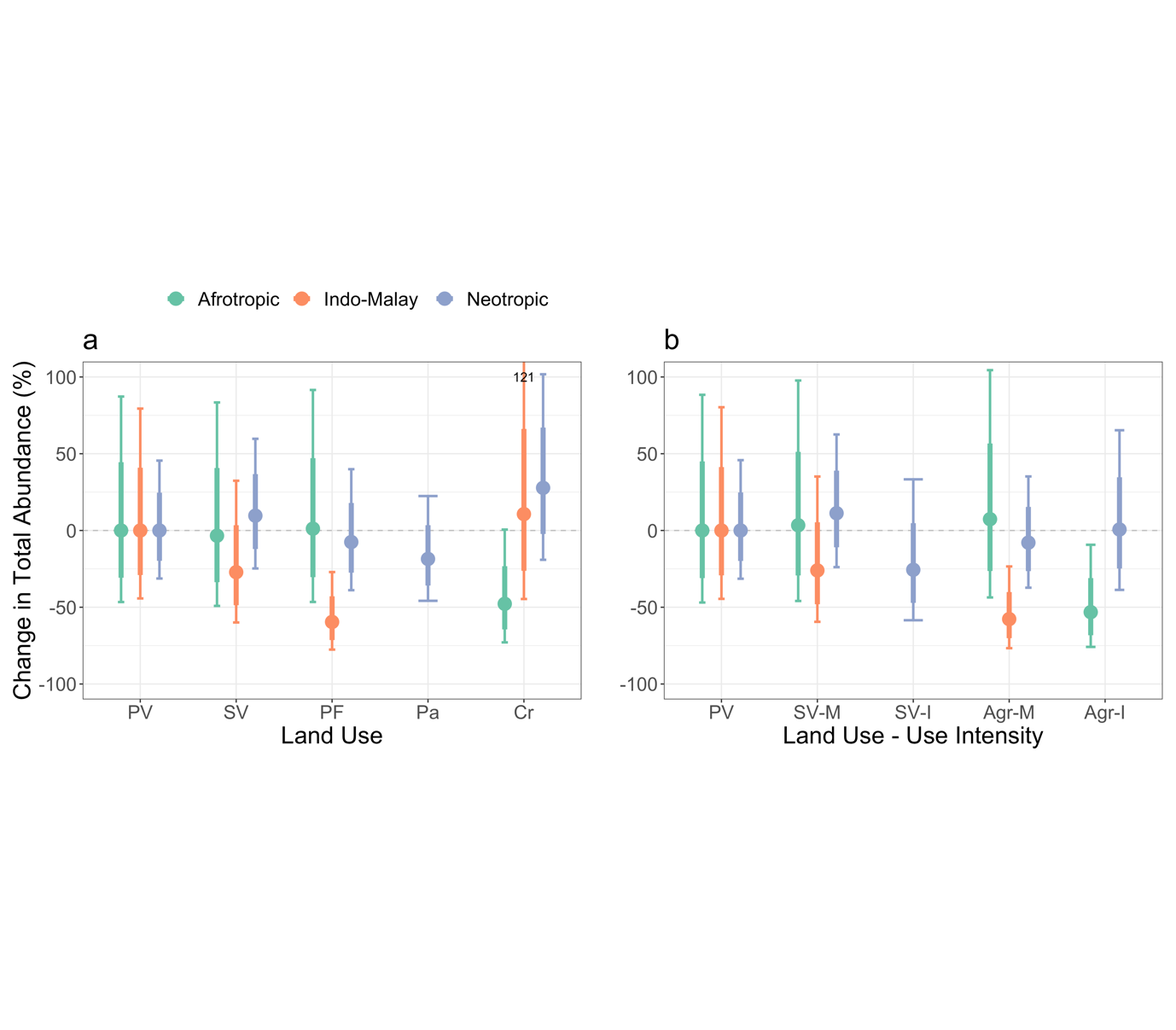

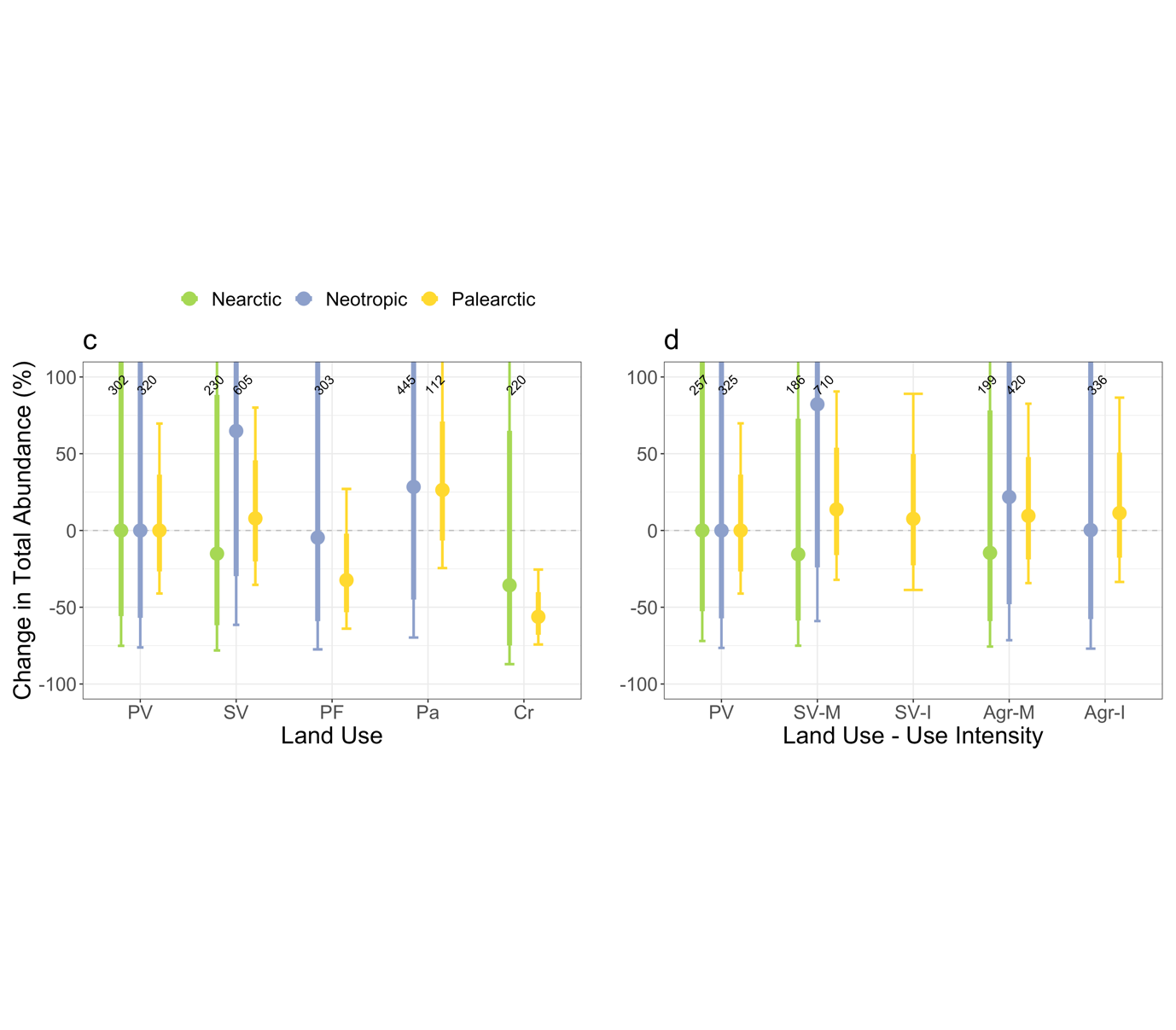

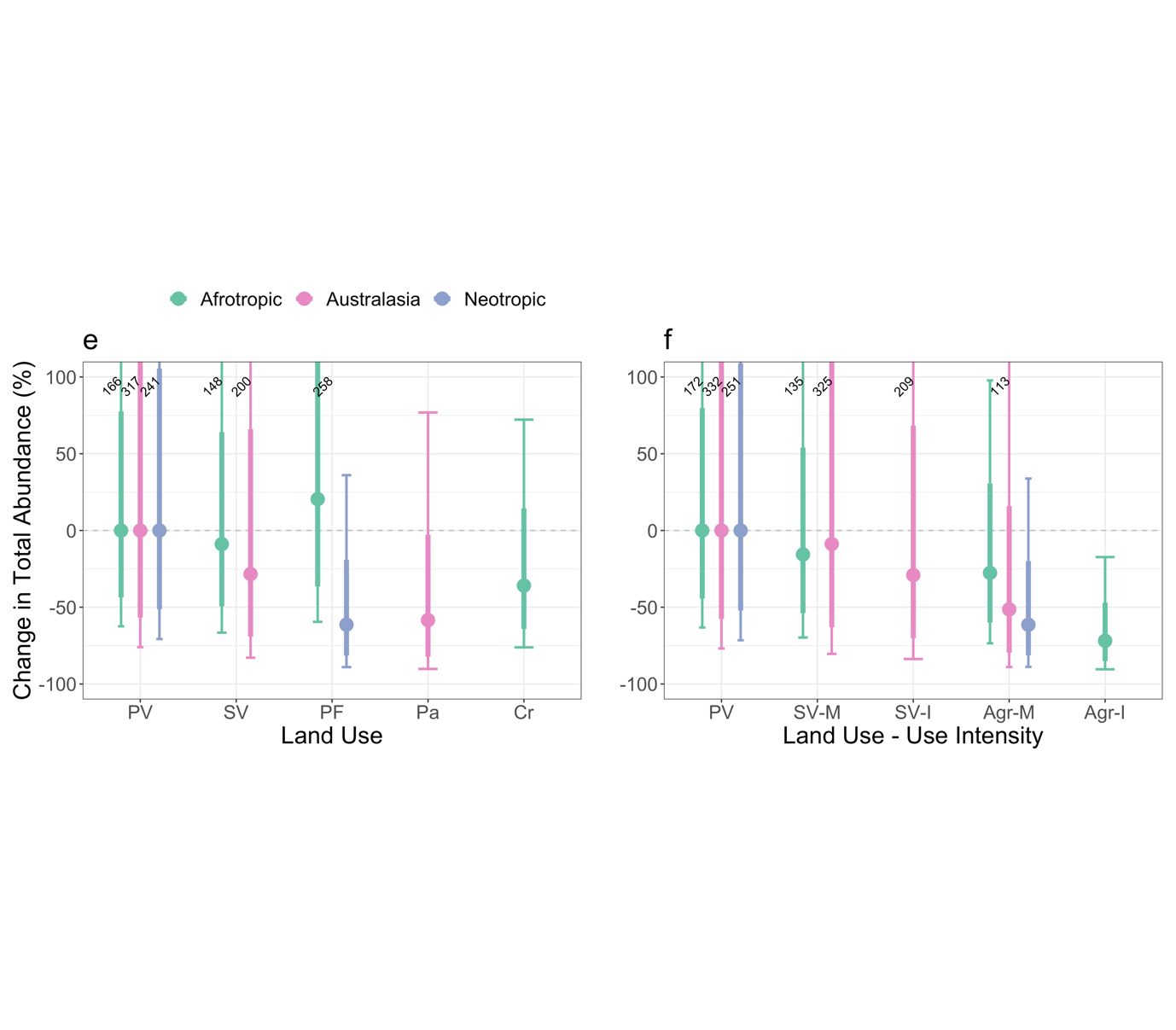
